# pr2-primers: an 18S rRNA primer database for protists

**DOI:** 10.1101/2021.01.04.425170

**Authors:** Daniel Vaulot, Stefan Geisen, Frédéric Mahé, David Bass

## Abstract

Metabarcoding of microbial eukaryotes (collectively known as *protists*) has developed tremendously in the last decade, almost uniquely relying on the 18S rRNA gene. As microbial eukaryotes are extremely diverse, many primers and primer pairs have been developed. To cover a relevant and representative fraction of the protist community in a given study system, a wise primer choice is needed as no primer pair can target all protists equally well. As such, a smart primer choice is very difficult even for experts and there are very few on-line resources available to list existing primers. We built a database listing 179 primers and 76 primer pairs that have been used for eukaryotic 18S rRNA metabarcoding. *In silico* performance of primer pairs was tested against two sequence databases: PR^2^ for eukaryotes and a subset of Silva for prokaryotes. This allowed to determine the taxonomic specificity of primer pairs, the location of mismatches as well as amplicon size. We developed a R-based web application that allows to browse the database, visualize the taxonomic distribution of the amplified sequences with the number of mismatches, and to test any user-defined primer set (https://app.pr2-primers.org). This tool will provide the basis for guided primer choices that will help a wide range of ecologists to implement protists as part of their investigations.

## Introduction

Microbes are key players in all Earth ecosystems. Among them are protists that encompass all unicellular or unicellular-colonial eukaryotes excluding some fungi. Protists perform a range of functions from photosynthesis to organic matter degradation. Although some eukaryotic groups such as unicellular algae (phytoplankton) have a long tradition of being studied as key players in marine primary production, the importance of protists in other processes and other environments has only been recently recognized, for example their role in nutrient cycling in soils or the complexity of symbiotic and parasitic relationships in marine waters (Geisen et al. 2018a; Worden et al. 2015). This absence of recognition stems in part from the inherent difficulties to identify them because of the lack of morphological features as well as of the difficulty to grow them in culture. In recent years the development of metabarcoding has provided new tools to study protist roles.

Metabarcoding is defined as the use of a specific marker gene to analyse the composition of natural communities in a specific environment (water, soil, animal gut, faeces, etc…). After DNA extraction, the gene is amplified using a pair of primers targeting one specific region, samples are labelled with molecular tags and the resulting DNA is sequenced using a high throughput technology, mostly Illumina currently. This approach was initially developed for prokaryotes (Sogin et al. 2006) and expanded steadily in recent years for protists. For prokaryotes, the gene most commonly used is the gene coding for the small sub-unit ribosomal RNA (SSU rRNA or 16S). SSU rRNA genes are composed of conserved and variable regions. Conserved regions can be used to design primers while variable region (V) can be used to assign taxonomy. In prokaryotes, the region targeted is very often V3/V4, although other regions have been suggested as providing better resolution (e.g. Bukin et al. 2019). For eukaryotes, earlier metabarcoding work done on microbial communities used the 18S rRNA gene (Amaral-Zettler et al. 2009; Stoeck et al. 2009), following up on what had been done using earlier cloning and Sanger sequencing approaches (López-García et al. 2001; Moon-van der Staay et al. 2001). Other genes, and in particular the mitochondrial cytochrome oxidase 1 gene (COI or cox1), have been used for Metazoa but their use is debated in particular because of the lack of universal primers (Andújar et al. 2018; Deagle et al. 2014) and the absence of this gene in lineages that have lost the mitochondrial genome (e.g. Yahalomi et al. 2020). For protists, the 18S rRNA gene appears to be most appropriate as a general marker (Pawlowski et al. 2012), although other genes such as rbcL (large unit of the RUBISCO) have been used for specific purposes such as targeting photosynthetic organisms (e.g. Pujari et al. 2019). Two variable regions of the 18S rRNA gene have been mostly targeted, V4 and V9: V4 is located in the second quarter of the 18S rRNA gene and V9 at the end of the 18S rRNA gene, near the ITS (internally transcribed spacer) region. Initially, the V4 region was favoured for 454 sequencing technology and V9 for Illumina which was then restricted to 2×75 bp. However with the development of the Illumina MiSeq (up to 2×300 bp), the V4 region is now preferred, in particular because it is longer, more variable, and better covered in reference databases (Pawlowski et al. 2012).

Primer selection is critical to obtain an accurate image of protist communities. Each primer (forward and reverse) must amplify the target community with minimal biases and the region amplified must be long enough to be taxonomically resolutive but preferably short enough to be fully sequenced by the chosen technology, although longer amplicons can be also be partially sequenced. With Illumina sequencing being now the most used technology, amplicon size must be about 50 bp smaller that the sum of the forward and reverse sequences (called R1 and R2) to allow enough overlap to reconstruct the complete amplicon: for example, the Illumina MiSeq 2×300 bp chemistry can sequence amplicons of up to 550 bp. A large diversity of primer and primer sets targeting the 18S rRNA gene have been developed over the years, although a few dominate in protist metabarcoding studies. Few resources are available that list eukaryotic 18S primers and primer pairs, provide information on their taxonomic specificity and allow to test new primer pairs. Most existing primer databases do not focus on protists. For example the primer database linked the Barcode of Life Data System web site Bold Systems (https://boldsystems.org/index.php/Public_Primer_PrimerSearch) is focusing on metazoans and Probebase (http://probebase.csb.univie.ac.at/node/8, Greuter et al. 2016) focuses on bacteria. A few programming tools have been developed to test primer set specificity, for example EcoPCR (Ficetola et al. 2010), a Python program, or R libraries such as PrimerMiner (Elbrecht and Leese 2017). Unfortunately, these tools need to be installed in a specific computing environment and require some background programming skills. Many existing online tools such as Probematch (https://rdp.cme.msu.edu/probematch/search.jsp) only allow testing primer sets against bacteria, archaea and fungi. Silva TestPrime (https://www.arb-silva.de/search/testprime) is the only tool that covers protists. It provides very detailed feedback on the taxonomy of amplified sequences, and the location of mismatches. Such detailed information comes at the expense of speed with a typical test needing a few minutes to run. Moreover the taxonomic annotation of the Silva database for protists is not optimal.

To fill this gap and to provide protist researchers with a usable tool, we constructed a database of primer and primer sets used for eukaryotic 18S rRNA metabarcoding. These primer sets were tested *in silico* against the PR^2^ database (Guillou et al. 2013) that contains more than 180,000 18S rRNA sequences with expert taxonomical annotation and a subset of the prokaryotic Silva database. We developed a R-based web application that allows to explore the database,d to visualize pre-computed *in silico* amplification results according to taxonomy (% of amplification, size of amplicons and location of mismatches), and to test any user-defined primer set.

## Material and Methods

18S rRNA gene primers (Table S1) and primer sets (Table S2) used in metabarcoding studies were collected from the literature. Primer sequences and primer sets (knowing that several primer sets may share at least one primer) were stored in a MySQL database. Primer sets were tested by performing *in silico* amplification of eukaryotic sequences stored in the PR^2^ reference database (Guillou et al. 2013) version 4.12.0 (https://github.com/pr2database/pr2database/releases/tag/v4.12.0). We also used a small subset of the Silva database version 132 provided by the mothur web site (https://mothur.org/wiki/silva_reference_files) to test whether prokaryotes were amplified. Sequences with ambiguities were discarded (any nucleotide that is not A, C, G or T). Sequences with length shorter than 1350 bp were not considered except for the V4 region for which this threshold was lowered to 1200 bp, since most sequences from PR^2^ contain the V4 region. In contrast, this limit was extended to 1650 for the V9 region and since many 18S rRNA do not cover the full V9 region, we only kept sequences that contained the canonical sequence GGATC[AT] which is located at the end of the V9 region, just before the start of the internally transcribed spacer 1 (ITS1). A R (R Development Core Team 2013) script using the *Biostrings* package (Pagès et al. 2020) was used to compute the number of mismatches to the forward and reverse primers allowing for a maximum of 2 mismatches for each primer using the function *matchPattern* with the following parameters: max.mismatch=2, min.mismatch=0, with.indels=FALSE, fixed=FALSE, algorithm=“auto”. We computed the position of mismatches using the *mismatch* function with parameter fixed=FALSE. A faster version of the script is also available that does not compute mismatch position using the vectorized form of the *matchPattern* function *(vmatchPattern*). The latter function is used in the Shiny application (see below) allowing users to test their own prime sets. The data were tabulated using the *dplyr* package and plotted using the *ggplot2* package (Wickham 2016). A R shiny application to interact with the database was developed using the following R packages: *shiny, shinyFeedback* and *shinycssloaders* (Sali and Attali 2020).

All scripts including those for the Shiny application are available at https://github.com/pr2database/pr2-primers.

## Results and Discussion

### Database of primers and primer sets

We have been able to recover from the literature a total of 102 general eukaryotic primers and 77 specific to some taxonomic groups (Tables 1 and S1, https://app.pr2-primers.org). Some of these primers were designed early on when researchers began to amplify and sequence the 18S rRNA gene (e.g. primers EukA and EukB, Medlin et al. 1988). More recently, researchers have been designing primer specific of some taxonomic groups mostly targeting the division-level (e.g. S19F and S15rF for Foraminifera, Morard et al. 2011) or class-level (e.g. primer PRYM03+3 for Prymnesiophyceae, Egge et al. 2013). Some primers are also designed to block specific taxa (e.g. 18SV1V2Block against the coral *Pocillopora damicornis*, Clerissi et al. 2018) to be used in combination with more general primers (18SV1V2F in this case) or to avoid amplification of some groups (e.g. EUK581-F and EUK1134-R which do not amplify Metazoa, Carnegie et al. 2003). These primers are used when looking at the eukaryotic microbiome of specific organisms (corals, oysters) to avoid amplification of host’s genes (Bass and del Campo 2020).

**Table 1:** Type of primers listed in the pr2-primers database. General primers target all eukaryotes and specific primers only certain taxonomic groups.

| direction | general | specific |
| --- | --- | --- |
| fwd | 52 | 42 |
| rev | 49 | 36 |

We identified a total of 76 primer sets that have been used in metabarcoding studies, mostly targeting protists (Table S2). Of these, most are general, i.e. not targeting specific groups. The distribution of these primer sets over the 18S rRNA gene is very heterogeneous, but the vast majority target the V4 region (Table 2 and Fig. 1). In contrast the number of primer sets targeting the other favoured metabarcoding region V9 is much lower. Most of the primer sets targeting a specific taxonomic group are located in the V4 region, and none are in the V9 region (Table 2). In terms of usage, the V4 region is much more popular (about 80% of published studies in marine systems, Lopes dos Santos et al. 2021), the three most commonly used primer sets being #8 (TAReuk454FWD1 and TAReukREV3, Stoeck et al. 2010), # 17 (E572F and E1009R, Comeau et al. 2011) and # 16 (TAReuk454FWD1 and V4 18S Next.Rev, Piredda et al. 2017), while for the V9 region the most popular sets are # 27 (1391F and EukB, Stoeck et al. 2010) and #28 (1380F and 1510R, Amaral-Zettler et al. 2009).

**Figure 1:**
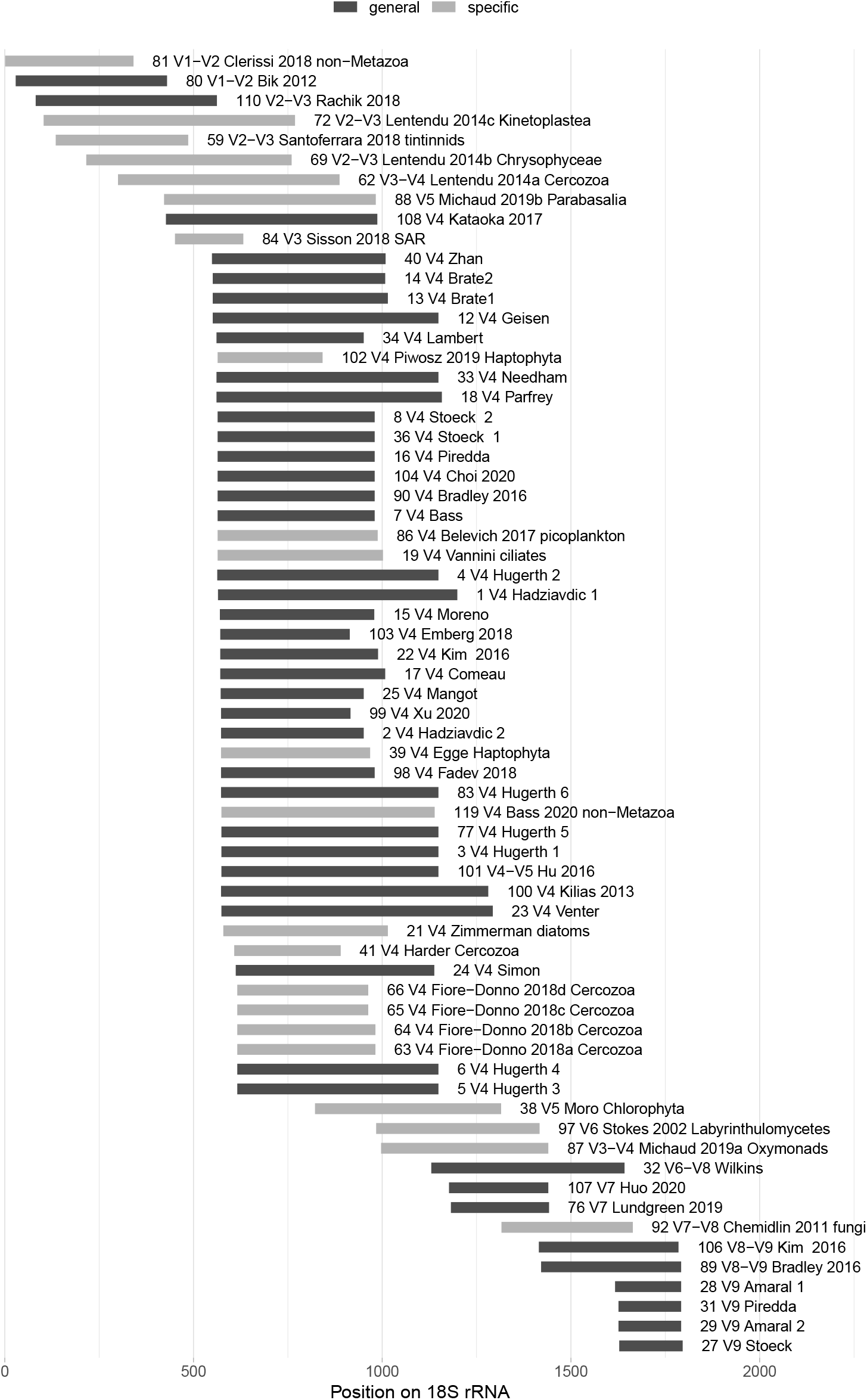
Position of the primer sets listed in the pr2-primers database along the 18S RNA gene relative to the sequence of the yeast *Saccharomyces cerevisiae* (FU970071). The label correspond to the primer set id, the 18S region amplified, its identification name and the specific group it eventually targets. Bar shading indicates whether the primer is general (black) or specific (grey) of a taxonomic group.

**Table 2:** Regions of the 18S rRNA gene targeted by the primer sets from the pr2-primers database.

| gene region | general | specific |
| --- | --- | --- |
| 37F-41F |  | 1 |
| V1-V2 | 1 | 1 |
| V2-V3 | 1 | 3 |
| V3 |  | 1 |
| V3-V4 |  | 2 |
| V4 | 32 | 13 |
| V4-V5 | 1 |  |
| V5 |  | 2 |
| V6 |  | 1 |
| V6-V8 | 1 |  |
| V7 | 2 |  |
| V7-V8 |  | 1 |
| V8-V9 | 2 |  |
| V9 | 4 |  |

### Testing primer sets by *in silico* matching

#### General primer sets

We used the PR^2^ database (Guillou et al. 2013) which currently contains about 180,000 18S rRNA sequences with a detailed taxonomic annotations to test all primer sets from the pr2-primers database. We also verified on a small set of sequences representative of the different prokaryotic groups whether these primers amplified bacteria or archaea. We only used long sequences (see Material and Methods) and allowed for a maximum of 2 mismatches on both forward and reverse primers, i.e. a maximum of 4 mismatches. For general primers, amplification success varied from 32 to more than 97% (Table 3, Fig. 2 and S1). In general, the reverse primer has a tendency to have more mismatches than the forward primer (Table 3). Primer sets targeting regions other than V4 or V9 do not perform as well in general (Fig. S1), although the best overall performance is for # 76 targeting the V7 region (F-1183 and R-1443, 97.1% of sequences amplified, Lundgreen et al. 2019). If we focus on the V4 and V9 regions (Fig. 2), the best performing primer sets overall are #6 (616*f and 1132r, 96.5%, Hugerth et al. 2014) and #29 (1389F and 1510R, 79.8%, Amaral-Zettler et al. 2009). The lower percentage observed for the V9 primers have to be interpreted with caution: many 18S reference sequences do not extend to the end of the V9 region and therefore will miss the signature of the reverse primer. To minimize this problem we retained for the analysis of V9 primer sets only sequences that contain the canonical signature GGATC[AT] located at the end of the V9 region. Despite performing well when allowing for 4 mismatches, some of these primer sets have at least one mismatch to PR^2^ sequences: for example primer set # 108 (545F and 1119R, Kataoka et al. 2017) amplifies only 7.9% of the sequences with zero mismatch. Another important consideration is the size of the amplicon. Since most metabarcoding studies are currently using Illumina sequencing technology, the maximum possible size to allow some overlap between the two R1 and R2 reads is about 550 bp (assuming that one uses the 2×300 bp sequencing kits), although smaller amplicons are preferable to allow more overlap. A sizeable fraction of the primer sets produce amplicon close to or larger than 600 bp (Fig. 2). The post sequencing analysis strategy in this case would be to only use one of the reads (R1 is in general less noisy) without trying to assemble R1 and R2 (as done in Lambert et al. 2019).

**Figure 2:**
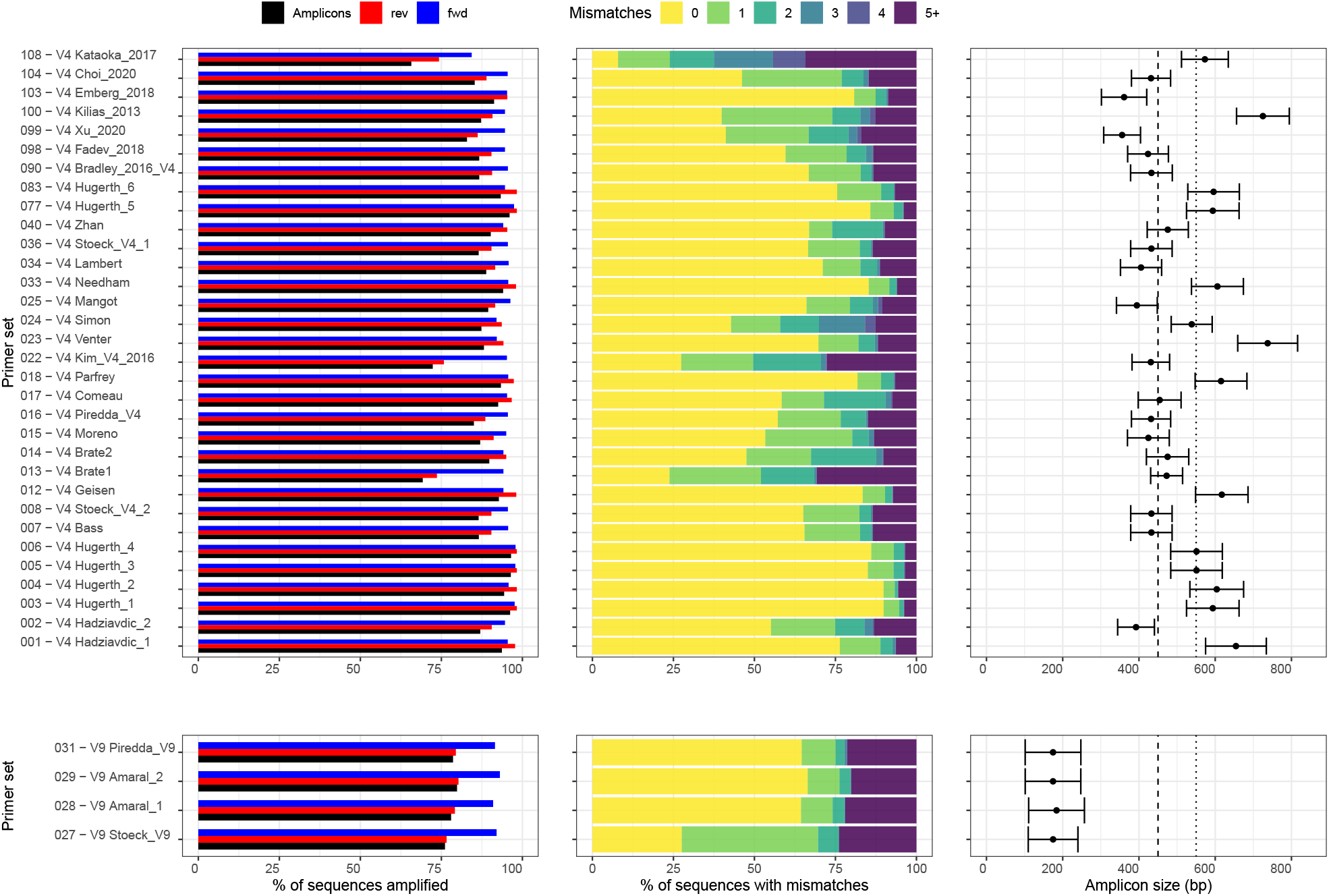
Evaluation of general primer sets (Table S2) targeting the V4 (top) and V9 (bottom) regions of the 18S rRNA gene against the PR^2^ reference database (version 4.12.0). Left panel. Percentage of reference sequences with at most 2 mismatches to either forward and reverse primer or to both primers, corresponding to the percentage of sequences amplified by the primer set. Central panel. Number of mismatches for each primer set. Right panel. Amplicon sizes targeted by different primer pairs. The vertical lines correspond to the lengths that can be covered by the most commonly used Illumina sequencers (dashed line: 2×250 base pairs; dotted line: 2×300 base pairs). Error bars represent the standard deviation.

**Table 3:** Overall characteristics of primer sets listed in the pr2-primers database.

|  | general | specific |
| --- | --- | --- |
| **forward primer** | % of sequences amplified |  |
| min | 36.4 | 0.0 |
| mean | 92.1 | 64.8 |
| max | 98.7 | 97.6 |
| **reverse primer** |  |  |
| min | 43.2 | 0.0 |
| mean | 89.9 | 32.4 |
| max | 98.6 | 96.1 |
| **both primers** |  |  |
| min | 30.0 | 0.0 |
| mean | 85.0 | 27.3 |
| max | 96.5 | 92.7 |
| **amplicon size** | bp |  |
| min | 174.6 | 184.0 |
| mean | 460.1 | 441.5 |
| max | 737.6 | 785.9 |

Another important consideration is whether amplification is similar across the whole eukaryotic taxonomic range. Taking as example the most used primer set targeting V4 (# 8, Fig. 3A) and looking at the amplification efficiency at the supergroup level, a significant fraction of Excavata and to a smaller extent of Rhizaria present at least 5 mismatches to this primer set (Fig. 3A top-left). Amplification is even more unlikely for sequences presenting mismatches with the forward primer because the mismatches are locate at the 3’ end of the primer (Fig. 3A top-right) which is the most unfavorable situation (mismatches at the 5’ end are better tolerated). The average size of the amplicon is also varying depending on the taxonomic group (Fig. 3B bottom). For example, Excavata have on average longer amplicons because of the presence of introns: amplicon size is then beyond the current range of Illumina sequencing and this may induce as well negative bias during PCR amplification (Geisen et al. 2015). For other groups such as Opisthokonta, although the average size is compatible with Illumina sequencing, there is a large number of outlier sequences with long amplicons. This will mean that taxa corresponding to these sequences (mostly Arthropoda) will be missed from surveys conducted with this primer set, although of course this is less critical when protists are targeted. The situation with V9 primer #27 (Fig. 3) is somewhat similar although there is less dissimilarity between the different supergroups. However for some supergroups, in particular Opisthokontha (Ascomycota) and Archae-plastida (Bangiophyceae), there is a number of outliers that will be missed by Illumina sequencing. Again these groups are less relevant when focusing on protists. When looking at all the general primer sets (Figure S2), some sets such as # 2, 25, and 110 appear to have more taxonomic biases than others. Overall Excavata constitute the supergroup that is most often discriminated against.

**Figure 3:**
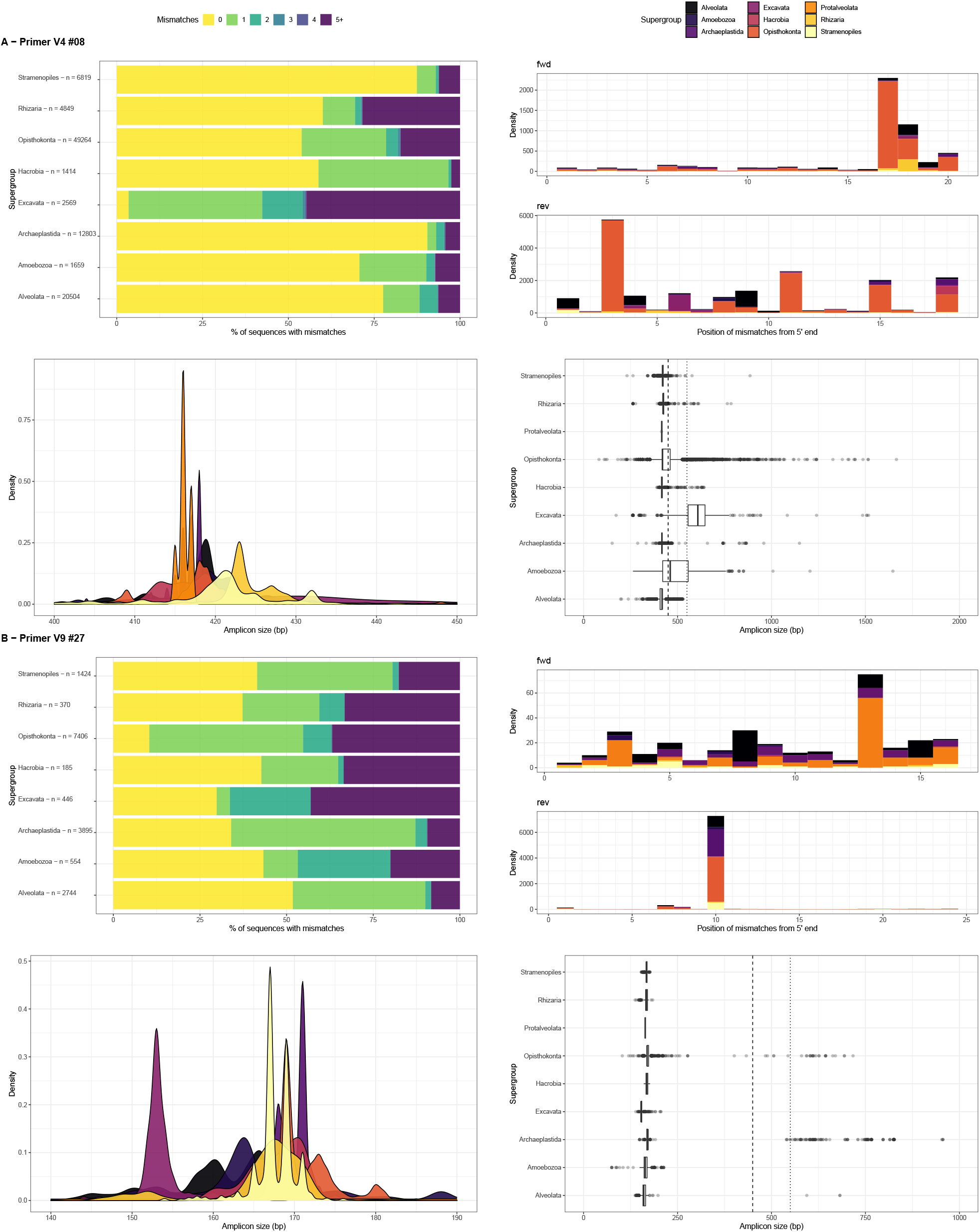
Example of analysis for two primer sets amplifying two regions of the 18S rRNA gene: V4 (primer set #8, A) and V9 (primer set #27, B). Top left. Percentage of sequences with a given number of mismatches. Top right. Position of the mismatches for different taxonomic supergroups on the forward and reverse primer counted from the 5’ end. Bottom left. Distribution of amplicon size for different supergroups. Bottom right. Box plots of amplicon size. Colors correspond to taxonomy (division). Hacrobia represents the sum of Haptophytes, Cryptophytes and Centrohelids.

Most primer sets will not amplify prokaryotes except primers such as set # 33 (515F and Univ 926R Needham and Fuhrman 2016) that were designed to amplify both bacteria and eukaryotes (Figs. S3 and S4). However some primers specific of eukaryotes such as # 4 (563f and 1132r, Hugerth et al. 2014) amplifies quite well prokaryotes. Interestingly, set # 12 (3NDf and 1132rmod, Geisen et al. 2018b) amplify only archaea but not bacteria. In most cases, it is the reverse primer which was discriminating against prokaryotes.

#### Specific primer sets

In order to access a deeper diversity within a given taxonomic group primers, primer sets have been developed with specific targets (Tables S1 and S2). Target levels are most often at the division (e.g. Haptophyta) and class levels (e.g. Chrysophyceae), although some sets are targeting supergroups (e.g. SAR #84). Some primer sets are extremely specific of their targets. One example is primer # 65 targeting Cercozoa (S616F Cerco and S947R Cerco, Fiore-Donno et al. 2018) that presents at least 5 mismatches to all other divisions (Fig. S5) and amplifies all Cercozoa groups. Primer # 38 targeting Chlorophyta (ChloroF and ChloroR, Moro et al. 2009) presents at least 5 mismatches to all other divisions (Fig. S5). However, it is does not amplifies all Chlorophyta as it misses picoplanktonic green algae such as Mamiellophyceae or Chloropicophyceae (Fig. S6). In contrast several primer sets claimed to be specific of a given group are in fact quite general. For example set # 87 which targets oxymonads (Oxy 18S-F and Oxy 18S-R Michaud et al. 2020) amplifies many other groups (Figs. S1 and S5). In this case, this is not critical since oxymonads only occur in termite guts and such primers will be used in this specific context. Primer set # 21 (D512for and D978rev, Zimmermann et al. 2011) which was designed to target diatoms would amplify actually most of the Ochrophyta (brown algae) classes but also some green algae (Fig. S6).

#### R Shiny application

We have developed a web site based on a R Shiny application (https://app.pr2-primers.org) that allows users to visualize and download the pr2-primers database, explore at different taxonomy levels the results of *in silico* amplification against the PR^2^ and Silva seed database for the primer sets from the database and test their own primer sets. The application is composed of 6 panels. The first panel (Fig. 4A) provides information on the database as well as link to report issues or new primers. The second and third panels (Fig. 4B) provide an interface to the two tables for primers and primers sets with the option of downloading it (item 1) and of revealing/hiding specific columns. The fourth and fifth panels allow to explore the *in silico* amplification of the primer sets from the database. The fourth panel present a synthesis of the results (similar to Fig. 2) while the fifth panel allows for a given primer set (item 2) to look at amplification properties within a taxonomic level from the kingdom to the class (item 3). The right panel shows general amplification characteristics (item 4), the location of the mismatches, the number of mismatches for each group and the distribution of the amplicon sizes (item 5). Finally the sixth panel allows users to provide a primer set and a maximum number of mismatches (item 6) to run an *in silico* amplification against PR^2^ and Silva seed databases. For the sake of speed, only the number of mismatches is provided but not the position of the mismatches as for the primer sets from the database. Global statistics on the amplification are provided which can be explored at different taxonomic levels (item 7). The R shiny application has been incorporated into a Docker container available at https://hub.docker.com/repository/docker/vaulot/pr2-primers.

**Figure 4:**
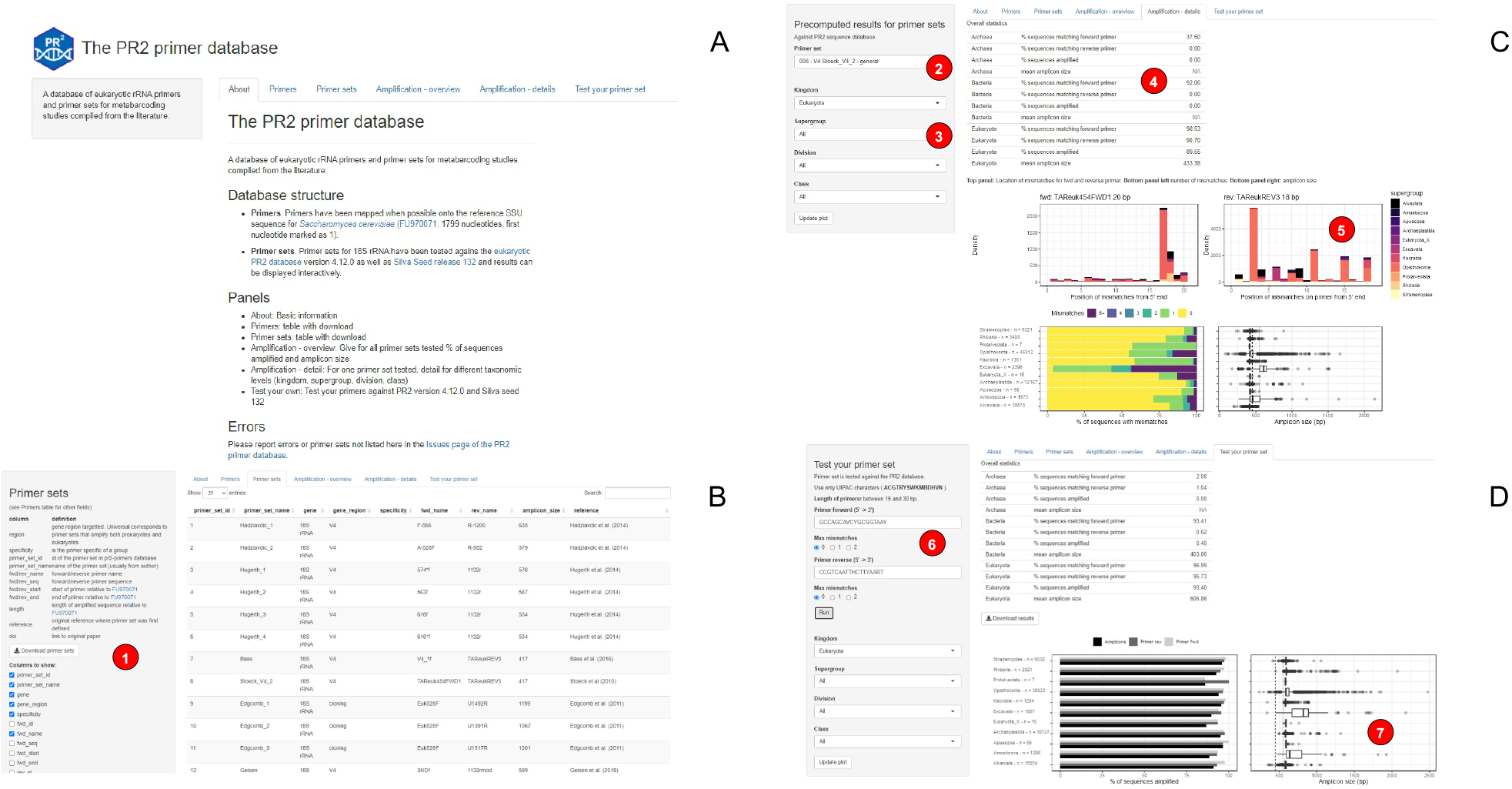
Shiny interface to the pr2-primers database. See text for details

## Conclusion

The combination of the pr2-primers database with the PR^2^ sequence database provides a very useful resource for protist metabarcoding. It will help researchers to select the most suitable primer pairs for both broadly-targeted surveys and studies focusing on target taxonomic groups, and to test and validate *in silico* novel primers. We emphasize that primer pairs must also be tested on reference culture material and natural samples as actual amplification may differ from *in silico* results. Hopefully this database will grow with time as novel primer pairs are developed and tested on samples from a range of environments. This will contribute to better design and comparability of microbiome analyses, inventories of protist diversity across environments, and increase our understanding of this functionally diverse and important group of organisms.

## Acknowledgements

We thank the ABIMS platform of the FR2424 (CNRS, Sorbonne Université) for bioinformatics resources.

## Author contributions statement

DV and SG conceived the study. DV, DB and FM scanned the literature for existing primers and primer sets. DV developed the database, the analysis scripts and the R shiny application. DV wrote the first draft of the paper and all co-authors edited and approved the final version.

## Additional information

### Competing interests

The authors declare no competing financial interests.

## Supplementary Material

**Table S1:**
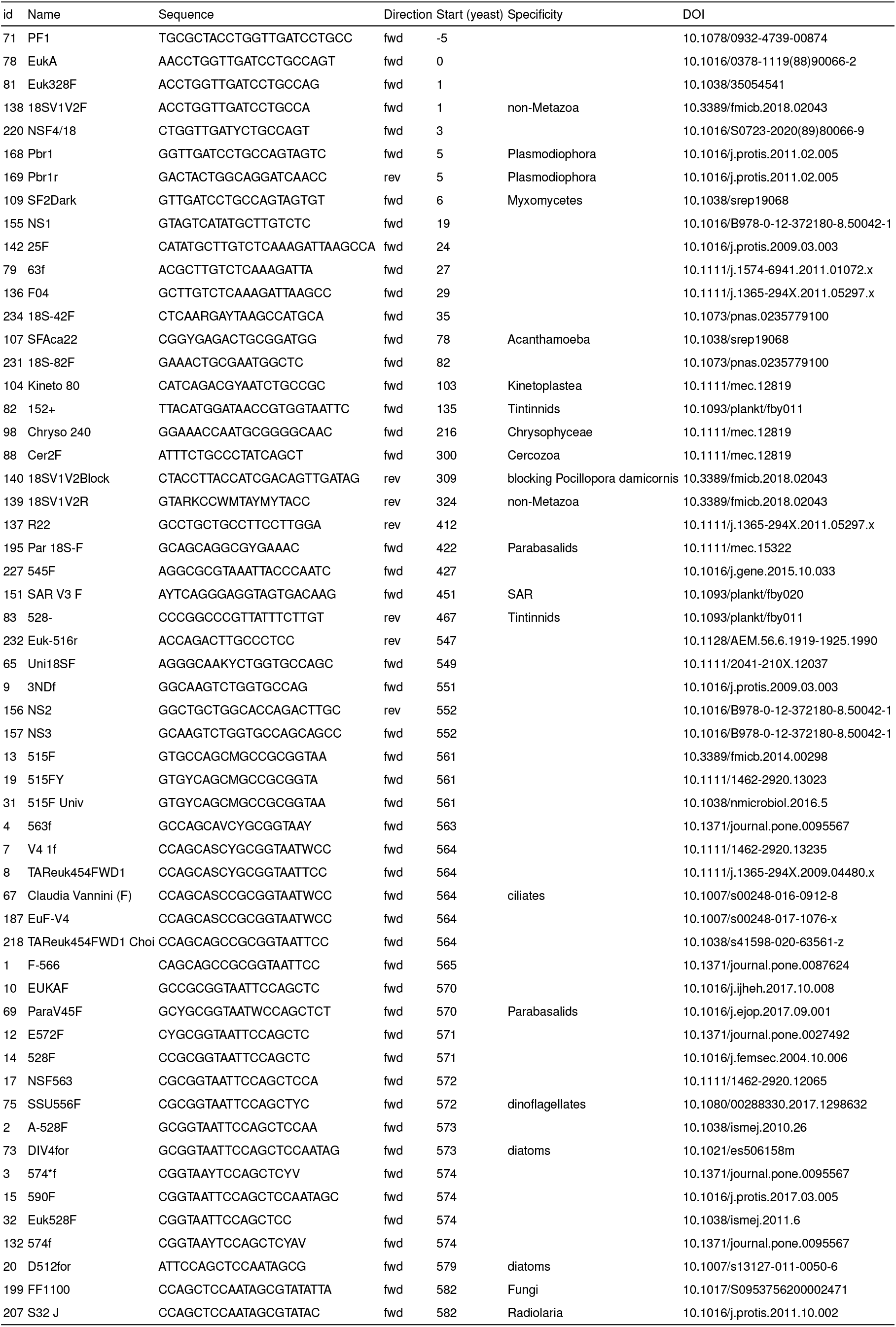
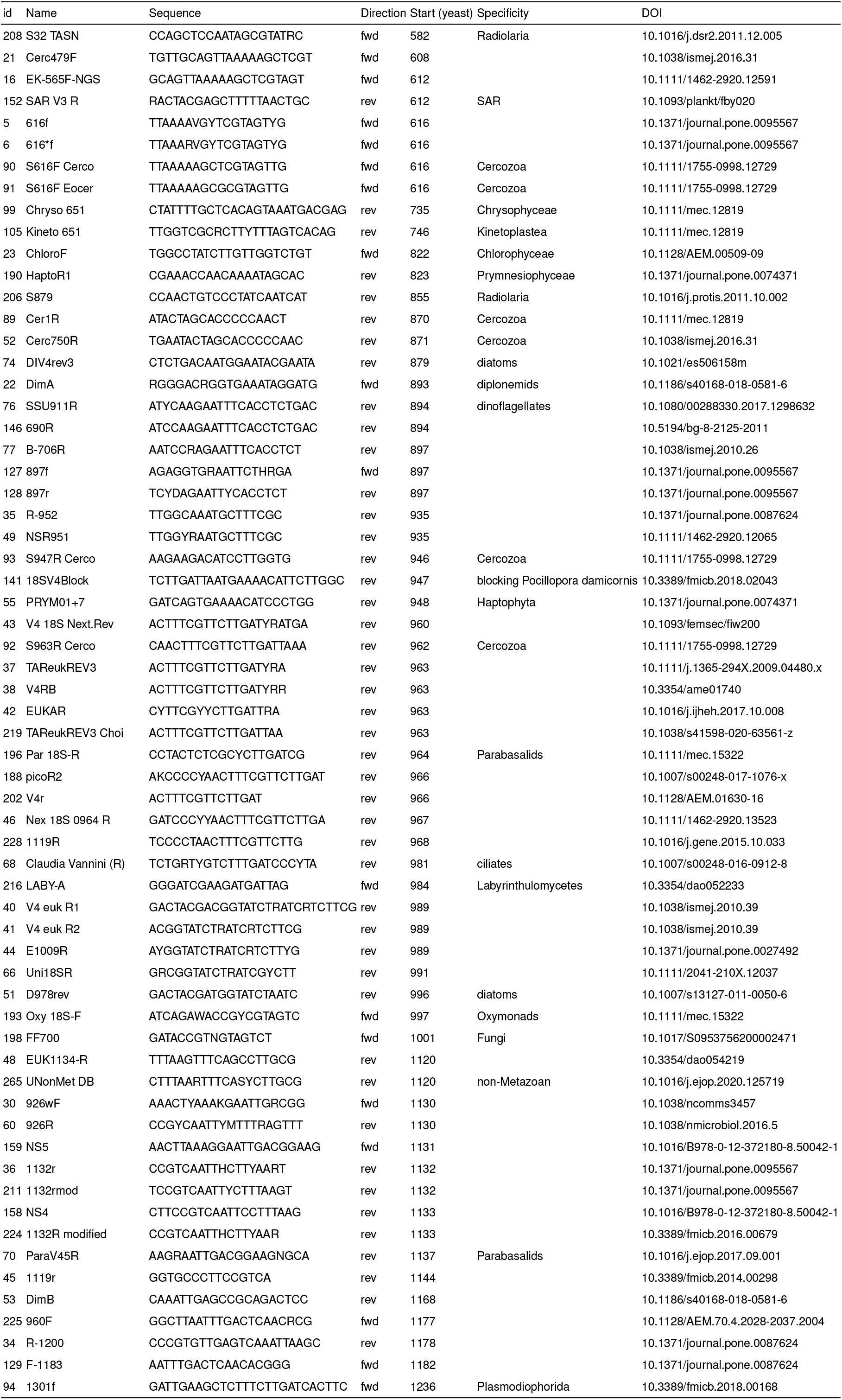
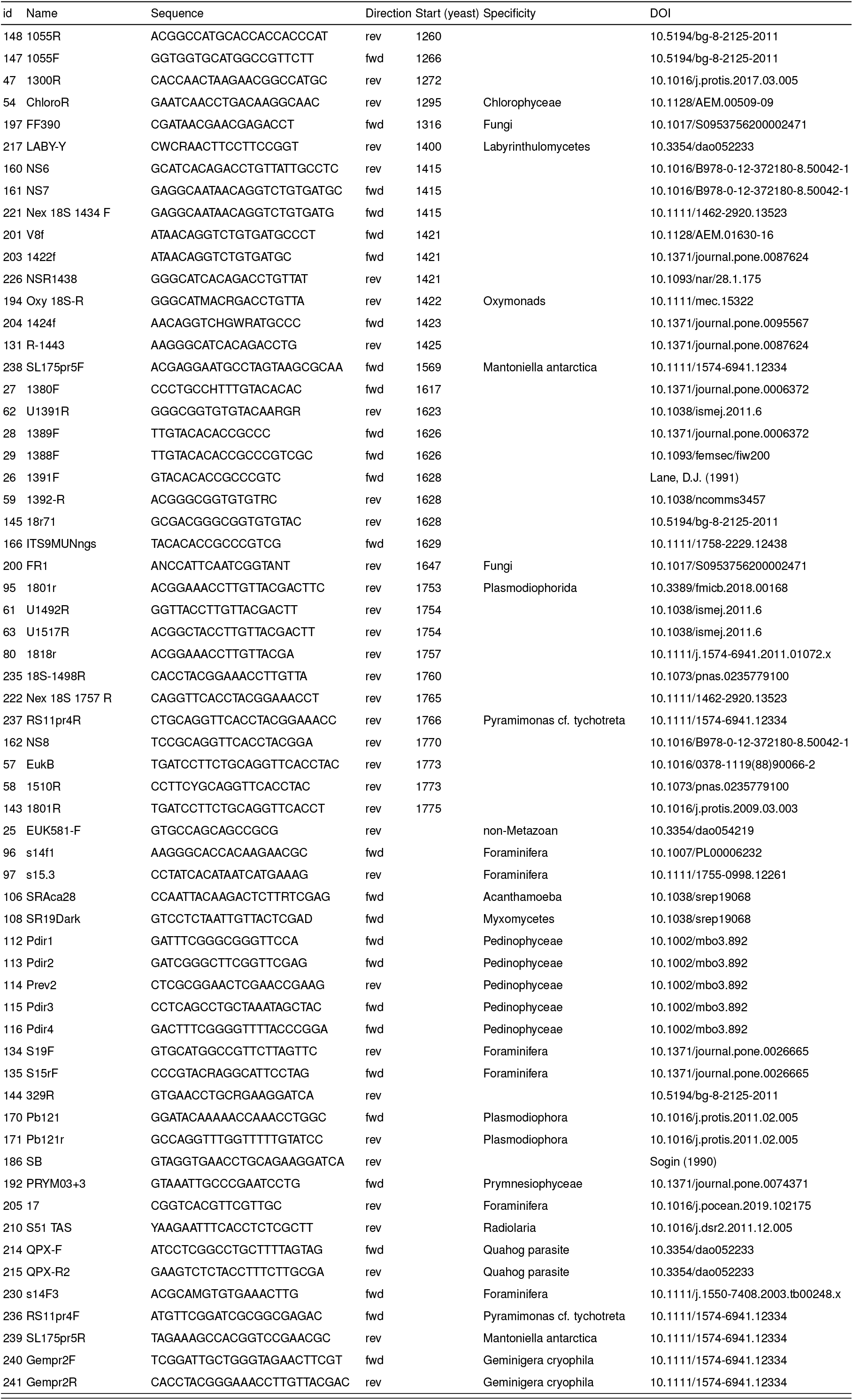
List of primers in the pr2-primers database ordered by start position relative to the sequence of the yeast extitSaccharomyces cerevisiae (FU970071).

**Table S2:**
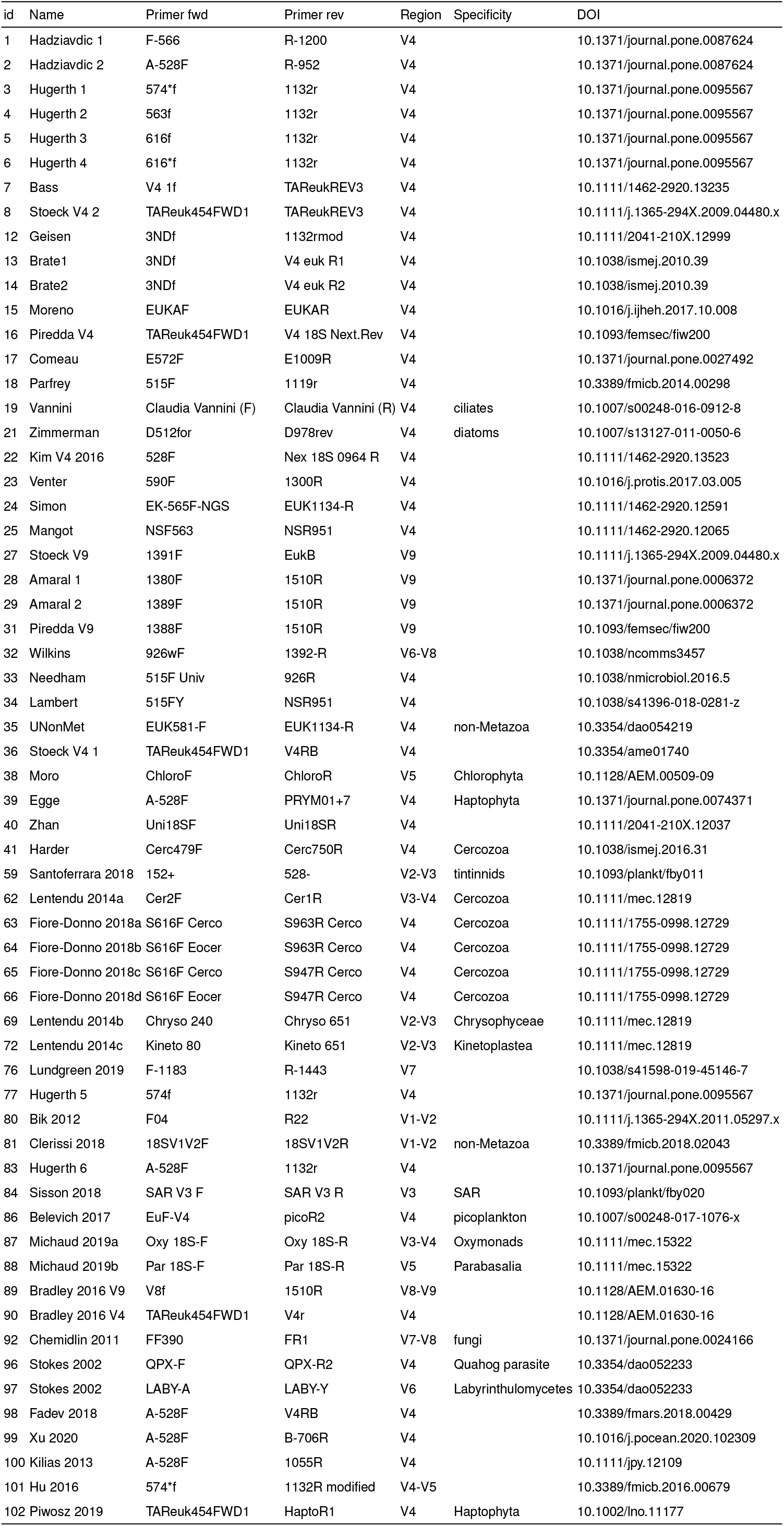
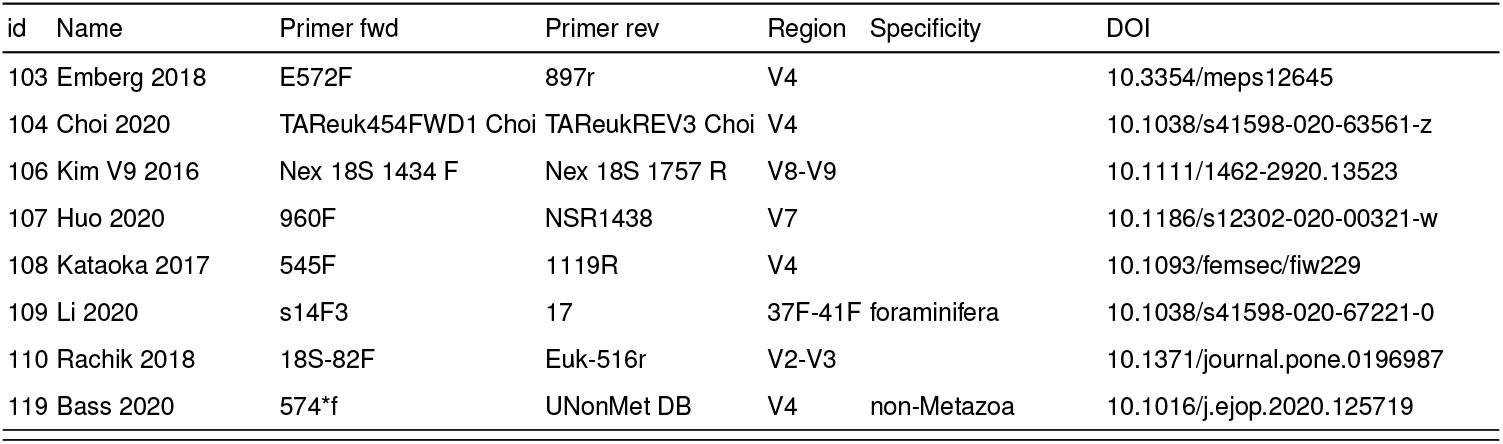
List of primer sets in the pr2-primers database.

**Figure S1:**
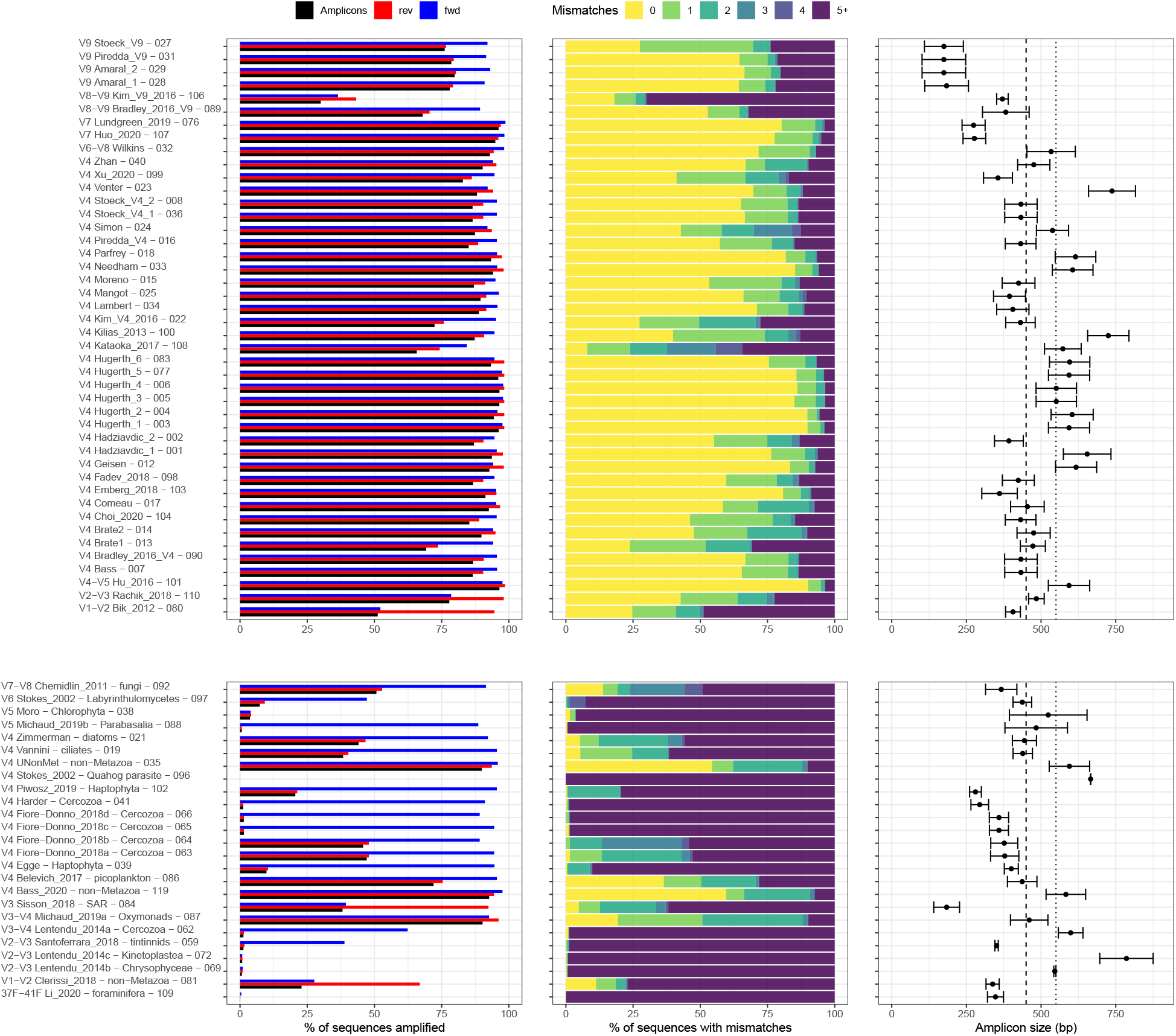
Evaluation of general (top) or specific (bottom) primer sets (Table S2) for the 18S rRNA gene against the PR^2^ reference database (version 4.12.0). See Fig. 2 for legend.

**Figure S2:**
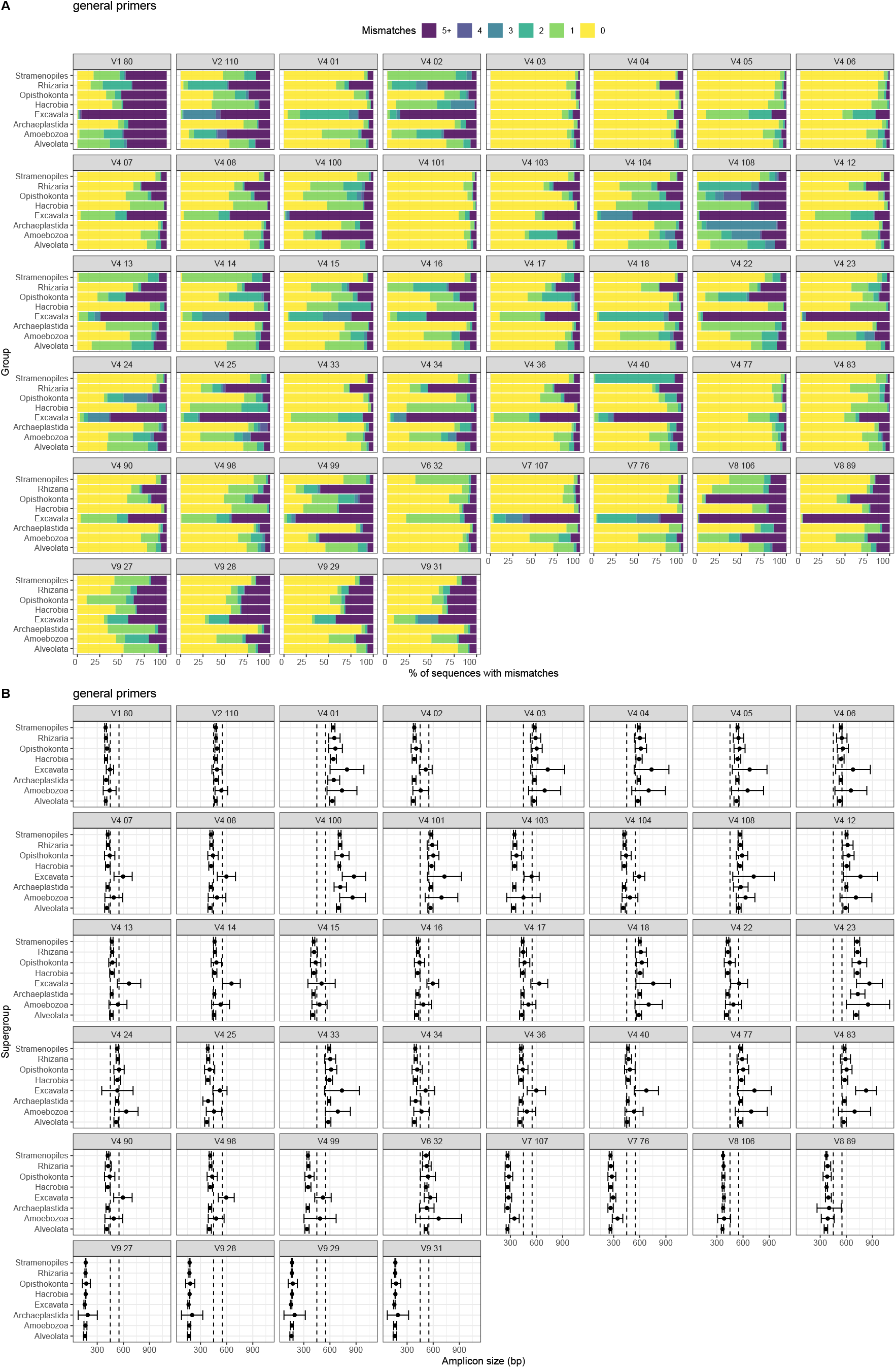
Number of mismatches (A) and amplicon size (mean ± SD, B) for general primer sets as a function of the supergroup.

**Figure S3:**
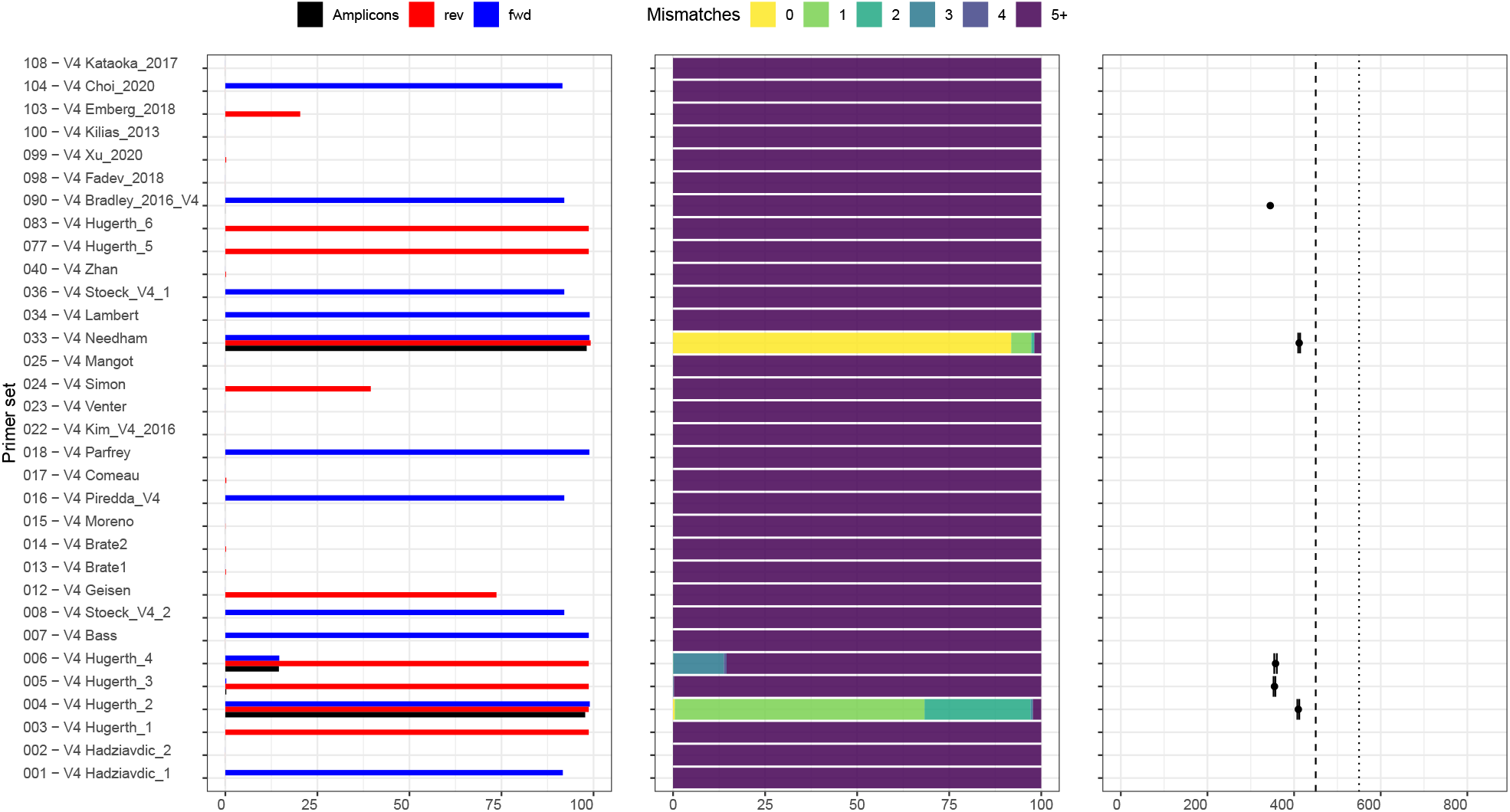
Evaluation of general primer sets (Table S2) targeting the V4 region of the 18S rRNA gene against bacterial 16S rRNA sequences from the Silva seed reference database (version 132). Legend as in Figure 2.

**Figure S4:**
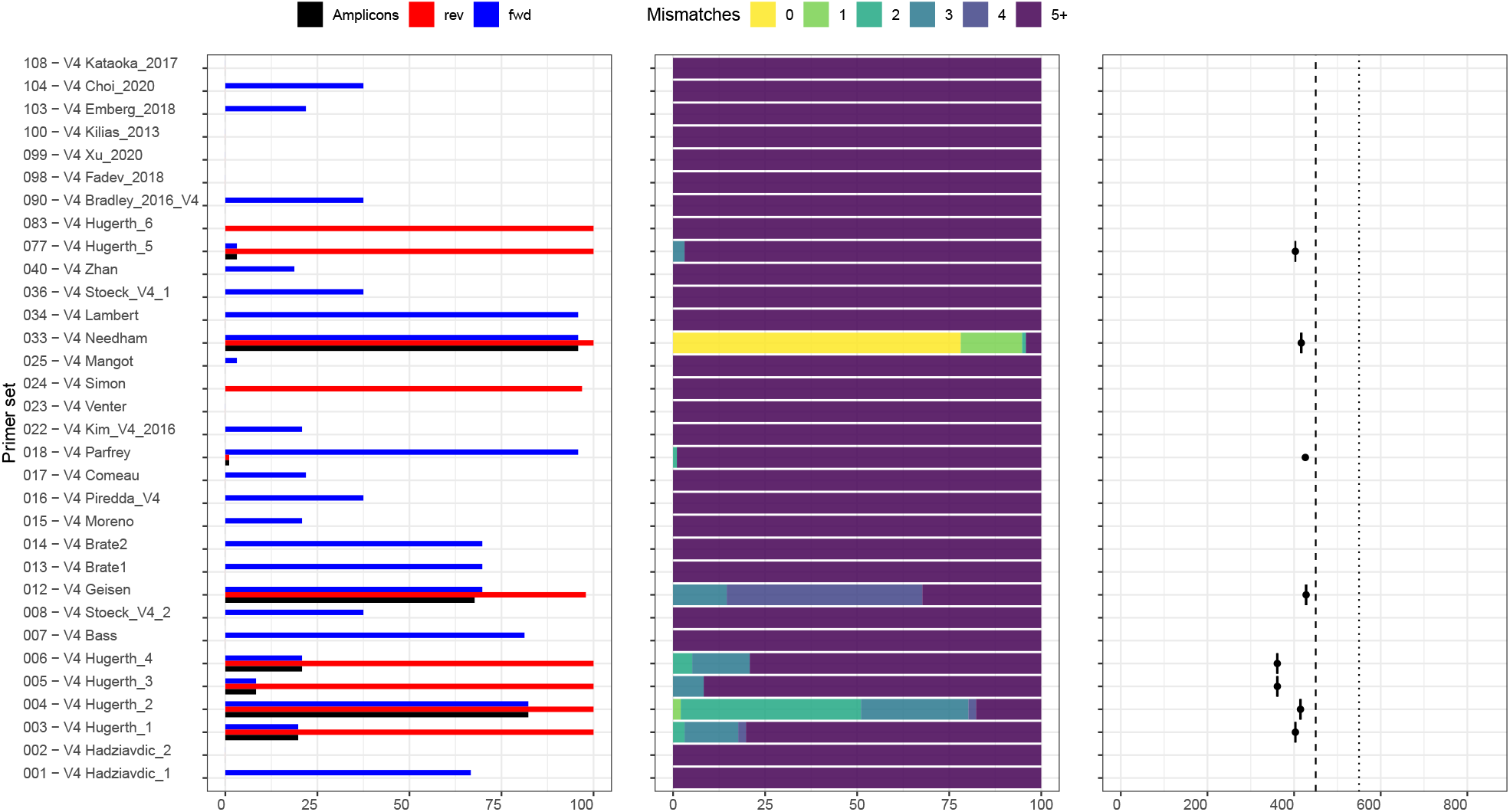
Evaluation of general primer sets (Table S2) targeting the V4 region of the 18S rRNA gene against archaeal 16S rRNA sequences from the Silva seed reference database (version 132). Legend as in Figure 2.

**Figure S5:**
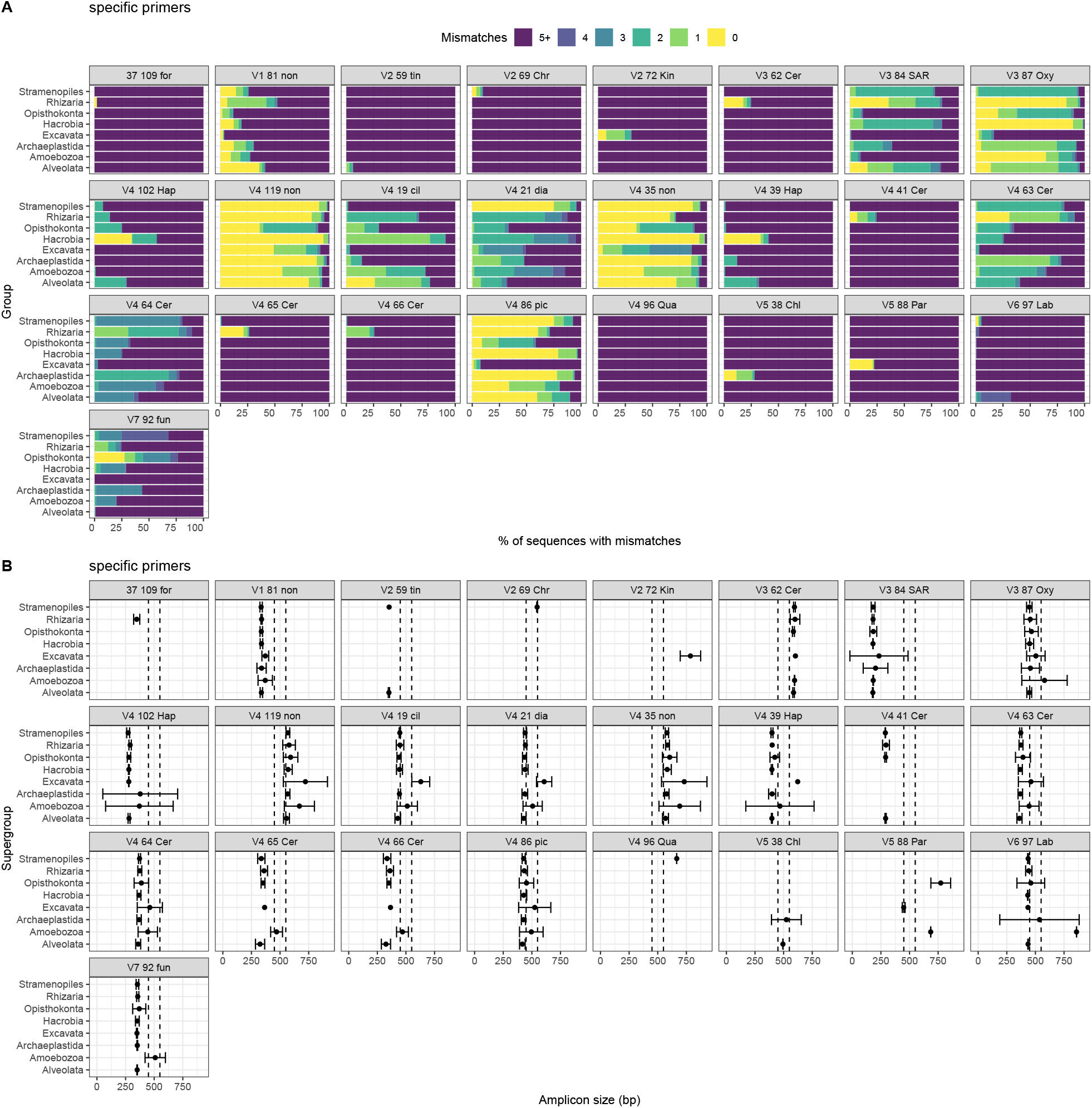
Number of mismatches (A) and amplicon size (mean ± SD, B) for specific primer sets as a function of the supergroup.

**Figure S6:**
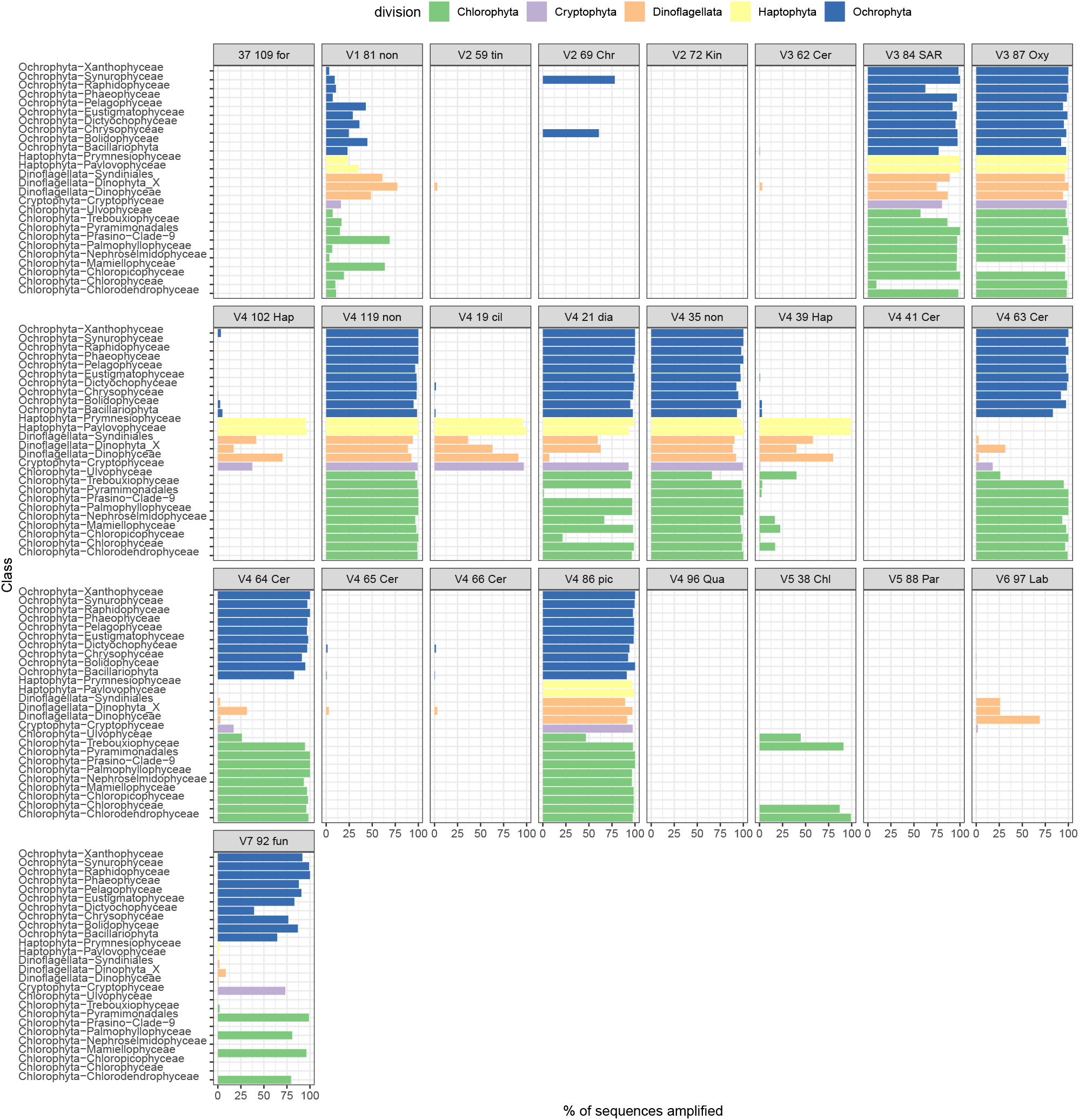
Percentage of sequences amplified with specific primer sets for different photosynthetic classes belonging to the Ochrophyta, Haptophyta, Dinoflagellata and Chlorophyta divisions.

## Notes

https://app.pr2-primers.org

https://github.com/pr2database/pr2-primers

### Competing Interest Statement

The authors have declared no competing interest.

## References cited

Amaral-Zettler, L. A., McCliment, E. A., Ducklow, H. W., & Huse, S. M. (2009). A Method for Studying Protistan Diversity Using Massively Parallel Sequencing of V9 Hypervariable Regions of Small-Subunit Ribosomal RNA Genes. PLoS ONE, 4, e6372. http://dx.doi.org/10.1371/journal.pone.0006372

Andújar, C., Arribas, P., Yu, D. W., Vogler, A. P., & Emerson, B. C. (2018). Why the COI barcode should be the community DNA metabarcode for the metazoa. Molecular Ecology, 27, 3968–3975. https://doi.org/10.1111/mec.14844

Bass, D., & del Campo, J. (2020). Microeukaryotes in animal and plant microbiomes: Ecologies of disease? European Journal of Protistology, 76, 125719. https://doi.org/10.1016/j.ejop.2020.125719

Bukin, Y. S., Galachyants, Y. P., Morozov, I. V., Bukin, S. V., Zakharenko, A. S., & Zemskaya, T. I. (2019). The effect of 16S rRNA region choice on bacterial community metabarcoding results. Scientific Data, 6, 190007. https://doi.org/10.1038/sdata.2019.7

Carnegie, R., Meyer, G., Blackbourn, J., Cochennec-Laureau, N., Berthe, F., & Bower, S. (2003). Molecular detection of the oyster parasite *Mikrocytos mackini*, and a preliminary phylogenetic analysis. Diseases of Aquatic Organisms, 54, 219–227. https://doi.org/10.3354/dao054219

Clerissi, C., Brunet, S., Vidal-Dupiol, J., Adjeroud, M., Lepage, P., Guillou, L., Escoubas, J.-M., & Toulza, E. (2018). Protists Within Corals: The Hidden Diversity. Frontiers in Microbiology, 9, 2043. https://doi.org/10.3389/fmicb.2018.02043

Comeau, A. M., Li, W. K. W., Tremblay, J.-É., Carmack, E. C., & Lovejoy, C. (2011). Arctic Ocean microbial community structure before and after the 2007 record sea ice minimum. PLoS ONE, 6, e27492. https://doi.org/10.1371/journal.pone.0027492

Deagle, B. E., Jarman, S. N., Coissac, E., Pompanon, F., & Taberlet, P. (2014). DNA metabarcoding and the cytochrome c oxidase subunit I marker: not a perfect match. Biology Letters, 10, 20140562. https://doi.org/10.1098/rsbl.2014.0562

Egge, E., Bittner, L., Andersen, T., Audic, S., de Vargas, C., & Edvardsen, B. (2013). 454 Pyrosequencing to Describe Microbial Eukaryotic Community Composition, Diversity and Relative Abundance: A Test for Marine Haptophytes. PLoS ONE, 8, e74371. https://doi.org/10.1371/journal.pone.0074371

Elbrecht, V., & Leese, F. (2017). PrimerMiner: an R package for development and *in silico* validation of DNA metabarcoding primers. Methods in Ecology and Evolution, 8, 622–626. https://doi.org/10.1111/2041-210X.12687

Ficetola, G., Coissac, E., Zundel, S., Riaz, T., Shehzad, W., Bessière, J., Taberlet, P., & Pompanon, F. (2010). An *In silico* approach for the evaluation of DNA barcodes. BMC Genomics, 11, 434. https://doi.org/10.1186/1471-2164-11-434

Fiore-Donno, A. M., Rixen, C., Rippin, M., Glaser, K., Samolov, E., Karsten, U., Becker, B., & Bonkowski, M. (2018). New barcoded primers for efficient retrieval of cercozoan sequences in high-throughput environmental diversity surveys, with emphasis on worldwide biological soil crusts. Molecular Ecology Resources, 18, 229–239. https://doi.org/10.1111/1755-0998.12729

Geisen, S., Laros, I., Vizcaíno, A., Bonkowski, M., & de Groot, G. A. (2015). Not all are free-living: high-throughput DNA metabarcoding reveals a diverse community of protists parasitizing soil metazoa. Molecular Ecology, 24, 4556–4569. https://doi.org/10.1111/mec.13238

Geisen, S., Mitchell, E. A., Adl, S., Bonkowski, M., Dunthorn, M., Ekelund, F., Fernández, L. D., Jousset, A., Krashevska, V., Singer, D., Spiegel, F. W., Walochnik, J., & Lara, E. (2018a). Soil protists: A fertile frontier in soil biology research. FEMS Microbiology Reviews, 42, 293–323. https://doi.org/10.1093/femsre/fuy006

Geisen, S., Snoek, L. B., ten Hooven, F. C., Duyts, H., Kostenko, O., Bloem, J., Martens, H., Quist, C. W., Helder, J. A., & der Putten, W. H. (2018b). Integrating quantitative morphological and qualitative molecular methods to analyse soil nematode community responses to plant range expansion. Methods in Ecology and Evolution, 9, 1366–1378. https://doi.org/10.1111/2041-210X.12999

Greuter, D., Loy, A., Horn, M., & Rattei, T. (2016). ProbeBase-an online resource for rRNA-targeted oligonucleotide probes and primers: New features 2016. Nucleic Acids Research, 44, D586–D589. https://doi.org/10.1093/nar/gkv1232

Guillou, L., Bachar, D., Audic, S., Bass, D., Berney, C., Bittner, L., Boutte, C., Burgaud, G., de Vargas, C., Decelle, J., del Campo, J., Dolan, J. R., Dunthorn, M., Edvardsen, B., Holzmann, M., Kooistra, W. H. C. F., Lara, E., Le Bescot, N., Logares, R., … Christen, R. (2013). The Protist Ribosomal Reference database (PR^2^): a catalog of unicellular eukaryote Small Sub-Unit rRNA sequences with curated taxonomy. Nucleic Acids Research, 41, D597–D604. https://doi.org/10.1093/nar/gks1160

Hugerth, L. W., Muller, E. E. L., Hu, Y. O. O., Lebrun, L. A. M., Roume, H., Lundin, D., Wilmes, P., & Andersson, A. F. (2014). Systematic Design of 18S rRNA Gene Primers for Determining Eukaryotic Diversity in Microbial Consortia. PLoS ONE, 9, e95567. http://dx.doi.org/10.1371/journal.pone.0095567

Kataoka, T., Yamaguchi, H., Sato, M., Watanabe, T., Taniuchi, Y., Kuwata, A., & Kawachi, M. (2017). Seasonal and geographical distribution of near-surface small photosynthetic eukaryotes in the western North Pacific determined by pyrosequencing of 18S rDNA. FEMS Microbiology Ecology, 93, fiw229. https://doi.org/10.1093/femsec/fiw229

Lambert, S., Tragin, M., Lozano, J.-C., Ghiglione, J.-F., Vaulot, D., Bouget, F.-Y., & Galand, P. E. (2019). Rhythmicity of coastal marine picoeukaryotes, bacteria and archaea despite irregular environmental perturbations. The ISME Journal, 13, 388–401. https://doi.org/10.1038/s41396-018-0281-z

Lopes dos Santos, A., Ong, D., Vaulot, D., Garczarek, L., Gérikas Ribero, C., Shi, X., & Gutiérrez-Rodríguez, A. (2021). Phytoplankton diversity and ecology through the lens of High Throughput sequencing technologies, In Advances in phytoplankton ecology: Applications of emerging technologies. Elsevier.

López-García, P., Rodriguez-Valera, F., Pedrós-Alió, C., & Moreira, D. (2001). Unexpected diversity of small eukaryotes in deep-sea Antarctic plankton. Nature, 409, 603–607.

Lundgreen, R. B. C., Jaspers, C., Traving, S. J., Ayala, D. J., Lombard, F., Grossart, H.-p., Nielsen, T. G., Munk, P., & Riemann, L. (2019). Eukaryotic and cyanobacterial communities associated with marine snow particles in the oligotrophic Sargasso Sea. Scientific Reports, 9, 8891. https://doi.org/10.1038/s41598-019-45146-7

Medlin, L., Elwood, H. J., Stickel, S., & Sogin, M. L. (1988). The characterization of enzymatically amplified eukaryotic 16S-like rRNA-coding regions. Gene, 71, 491–499. https://doi.org/10.1016/0378-1119(88)90066-2

Michaud, C., Hervé, V., Dupont, S., Dubreuil, G., Bézier, A. M., Meunier, J., Brune, A., & Dedeine, F. (2020). Efficient but occasionally imperfect vertical transmission of gut mutualistic protists in a wood-feeding termite. Molecular Ecology, 29, 308–324. https://doi.org/10.1111/mec.15322

Moon-van der Staay, S. Y., De Wachter, R., & Vaulot, D. (2001). Oceanic 18S rDNA sequences from picoplankton reveal unsuspected eukaryotic diversity. Nature, 409, 607–610. https://doi.org/10.1038/35054541

Morard, R., Quillévéré, F., Douady, C. J., de Vargas, C., de Garidel-Thoron, T., & Escarguel, G. (2011). Worldwide Genotyping in the Planktonic Foraminifer *Globoconella inflata:* Implications for Life History and Paleoceanography. PLoS ONE, 6, e26665. https://doi.org/10.1371/journal.pone.0026665

Moro, C. V., Crouzet, O., Rasconi, S., Thouvenot, A., Coffe, G., Batisson, I., & Bohatier, J. (2009). New design strategy for development of specific primer sets for PCR-based detection of Chlorophyceae and Bacillariophyceae in environmental samples. Applied and Environmental Microbiology, 75, 5729–5733. https://doi.org/10.1128/AEM.00509-09

Needham, D. M., & Fuhrman, J. A. (2016). Pronounced daily succession of phytoplankton, archaea and bacteria following a spring bloom. Nature Microbiology, 1, 16005. https://doi.org/10.1038/nmicrobiol.2016.5

Pagès, H., Aboyoun, P., Gentleman, R., & DebRoy, S. (2020). Biostrings: Efficient manipulation of biological strings [R package version 2.56.0]. R package version 2.56.0.

Pawlowski, J., Audic, S., Adl, S., Bass, D., Belbahri, L., Berney, C., Bowser, S. S., Cepicka, I., Decelle, J., Dunthorn, M., Fiore-Donno, A. M., Gile, G. H., Holzmann, M., Jahn, R., Jirků, M., Keeling, P. J., Kostka, M., Kudryavtsev, A., Lara, E., … de Vargas, C. (2012). CBOL Protist Working Group: Barcoding Eukaryotic Richness beyond the Animal, Plant, and Fungal Kingdoms. PLoS Biology, 10, e1001419. http://dx.doi.org/10.1371/journal.pbio.1001419

Piredda, R., Tomasino, M. P., D’Erchia, A. M., Manzari, C., Pesole, G., Montresor, M., Kooistra, W. H. C. F., Sarno, D., & Zingone, A. (2017). Diversity and temporal patterns of planktonic protist assemblages at a Mediterranean Long Term Ecological Research site. FEMS Microbiology Ecology, 93, fiw200. https://doi.org/10.1093/femsec/fiw200

Pujari, L., Wu, C., Kan, J., Li, N., Wang, X., Zhang, G., Shang, X., Wang, M., Zhou, C., & Sun, J. (2019). Diversity and spatial distribution of chromophytic phytoplankton in the bay of bengal revealed by RuBisCO Genes (rbcL). Frontiers in Microbiology, 10, 1–17. https://doi.org/10.3389/fmicb.2019.01501

R Development Core Team. (2013). R: A Language and Environment for Statistical Computing. https://doi.org/10.1007/978-3-540-74686-7

Sali, A., & Attali, D. (2020). Shinycssloaders: Add loading animations to a ’shiny’ output while it’s recalculating [R package version 1.0.0]. R package version 1.0.0. https://CRAN.R-project.org/package=shinycssloaders

Sogin, M. L., Morrison, H. G., Huber, J. A., Welch, D. M., Huse, S. M., Neal, P. R., Arrieta, J. M., & Herndl, G. J. (2006). Microbial diversity in the deep sea and the underexplored “rare biosphere”. Proceedings of the National Academy of Sciences of the United States of America, 103, 12115–12120.

Stoeck, T., Bass, D., Nebel, M., Christen, R., Jones, M. D. M., Breiner, H. W., & Richards, T. A. (2010). Multiple marker parallel tag environmental DNA sequencing reveals a highly complex eukaryotic community in marine anoxic water. Molecular Ecology, 19, 21–31. https://doi.org/10.1111/j.1365-294X.2009.04480.x

Stoeck, T., Behnke, A., Christen, R., Amaral-Zettler, L., Rodriguez-Mora, M. J., Chistoserdov, A., Orsi, W., & Edgcomb, V. P. (2009). Massively parallel tag sequencing reveals the complexity of anaerobic marine protistan communities. BMC Biology, 7, 72. https://doi.org/10.1186/1741-7007-7-72

Wickham, H. (2016). Ggplot2: Elegant graphics for data analysis. Springer International Publishing. https://books.google.com.sg/books?id=RTMFswEACAAJ

Worden, A. Z., Follows, M. J., Giovannoni, S. J., Wilken, S., Zimmerman, A. E., & Keeling, P. J. (2015). Rethinking the marine carbon cycle: Factoring in the multifarious lifestyles of microbes. Science, 347, 1257594. https://doi.org/10.1126/science.1257594

Yahalomi, D., Atkinson, S. D., Neuhof, M., Chang, E. S., Philippe, H., Cartwright, P., Bartholomew, J. L., & Huchon, D. (2020). A cnidarian parasite of salmon (Myxozoa: *Henneguya*) lacks a mitochondrial genome. Proceedings of the National Academy of Sciences, 117, 5358–5363. https://doi.org/10.1073/pnas.1909907117

Zimmermann, J., Jahn, R., & Gemeinholzer, B. (2011). Barcoding diatoms: Evaluation of the V4 subregion on the 18S rRNA gene, including new primers and protocols. Organisms Diversity and Evolution, 11, 173–192. https://doi.org/10.1007/s13127-011-0050-6

